# Towards A Wireless Image Sensor for Real-Time Fluorescence Microscopy in Cancer Therapy

**DOI:** 10.1101/2023.12.03.569779

**Authors:** Rozhan Rabbani, Hossein Najafiaghdam, Micah Roschelle, Efthymios Philip Papageorgiou, Biqi Rebekah Zhao, Mohammad Meraj Ghanbari, Rikky Muller, Vladimir Stojanovic, Mekhail Anwar

**Affiliations:** Department of Electrical Engineering and Computer Sciences, University of California at Berkeley, Berkeley, CA 94720 USA; Chan Zuckerberg Biohub, San Francisco, CA 94158 USA; Department of Radiation Oncology, University of California, San Francisco, CA 94158 USA

**Keywords:** Biomedical implants, cancer detection, capacitive trans-impedance amplifier (CTIA), energy harvesting, fluorescence imaging, image sensor, piezoceramic, ultrasound backscattering

## Abstract

We present a mm-sized, ultrasonically powered lensless CMOS image sensor as a progress towards wireless fluorescence microscopy. Access to biological information within the tissue has the potential to provide insights guiding diagnosis and treatment across numerous medical conditions including cancer therapy. This information, in conjunction with current clinical imaging techniques that have limitations in obtaining images continuously and lack wireless compatibility, can improve continual detection of multicell clusters deep within tissue. The proposed platform incorporates a 2.4×4.7 mm^2^ integrated circuit (IC) fabricated in TSMC 0.18 μm, a micro laser diode (μLD), a single piezoceramic and off-chip storage capacitors. The IC consists of a 36×40 array of capacitive trans-impedance amplifier-based pixels, wireless power management and communication via ultrasound and a laser driver all controlled by a Finite State Machine. The piezoceramic harvests energy from the acoustic waves at a depth of 2 cm to power up the IC and transfer 11.5 kbits/frame via backscattering. During *Charge-Up*, the off-chip capacitor stores charge to later supply a high-power 78 mW μLD during *Imaging*. Proof of concept of the imaging front end is shown by imaging distributions of CD8 T-cells, an indicator of the immune response to cancer, *ex vivo*, in the lymph nodes of a functional immune system (BL6 mice) against colorectal cancer consistent with the results of a fluorescence microscope. The overall system performance is verified by detecting 140 μm features on a USAF resolution target with 32 ms exposure time and 389 ms ultrasound backscattering.

## I. INTRODUCTION

**C**ONTINUOUS access to *in vivo* information through implantable biomedical sensors can provide insights for diagnosis and personalized treatment guidance based on real-time feedback from the patient’s own tissue [1]–[5]. These platforms are poised to address a critical challenge across medicine which is the wide heterogeneity in therapeutic response across patients. Therefore, an imaging method to visualize the real-time response to therapy to enable personalized medicine is needed [6]. Monitoring real-time changes deep inside the body with sufficient resolution is critical for understanding the dynamics of biological processes. For instance, access to *in vivo* imaging information is critical in cancer therapy and particularly immunotherapy, an effective treatment that unlocks the patient’s immune system to find and attack cancer cells [7]. Multiple clinical trials have demonstrated an improvement in overall survival with immunotherapy [8], [9]. However, less than 20-30% of the patients respond [10] and the therapy is not without risks including autoimmune side effects [11] and financial costs (∼$100,000 per patient [12]). Most importantly, time spent on ineffective immunotherapy closes the window for finding an effective cure. Given the low response rates to immunotherapy, rapid recognition of nonresponders and adaptation of therapy based on very early response assessment is critical for more effective, personalized therapeutic regimens. A key goal in evaluating immunotherapy is to image the complex immune response and capture the change in tumor state and distribution of clusters of cells that govern the response [13]. Visualizing tumor changes at the earliest possible time point, may reveal response or resistance far earlier than conventional methods. Therefore, an imaging system for cancer immunotherapy must be capable of imaging tumor state changes at time intervals frequent enough to capture cell cluster motion (∼minutes to hours), over long time periods (days to months).

Currently, these factors are only visible when looking at the tissue under a microscope and only with a biopsy. However, repeated biopsies, often of sites deep within the patient, are impractical due to morbidity, cost, and logistics. In clinical practice, the state of the art for monitoring immunotherapy response is to use clinical imaging technologies: computed tomography (CT), magnetic resonance imaging (MRI), positron emission tomography (PET) and multimodal techniques (CT/PET, PET/MRI, etc.) [14], [15]. Guidelines such as iRECIST [16] mandate volumetric changes of 20% in the tumor size to confirm disease progression, so even for a small tumor with a major dimension of 1-3 cm, this equates to a minimum change of 0.2 cm^3^. Given a cell is ∼10 μm, this equates to a minimum change of 200 million cells, which can take months to manifest [16]. While MRI and CT are invaluable in detecting anatomic changes, for non-responders, the window for rapid therapeutic adaptation is lost due to the inability to detect the early response in real time. Additionally, molecular PET-based imaging lacks the resolution necessary to monitor cancer state changes. Moreover, hospital-based imagers preclude serial imaging due to the logistics and cost associated with repeated imaging spaced only by hours or days. Without continuous monitoring, conventional techniques are restricted to snapshots unable to capture the dynamics of the key biological phenomena. Therefore, an additional method in conjunction with clinical imaging of the tumor response is needed.

Optical microscopy, on the other hand, enables high-resolution, multiplexed imaging providing key information regarding disease progression and treatment efficacy [17]. Fluorescence microscopy is an optical imaging method that relies on fluorophores that absorb light and emit light of a longer wavelength (i.e., lower energy). Fluorescence microscopy with the systemic injection of fluorescently-tagged cell-specific antibodies or small molecules enables 1) real-time monitoring with 2) sufficient resolution for capturing tumor stage changes and 3) visualizing multiple cell types simultaneously, which is not viable with other imaging modalities. With appropriate optical filters such as multi-bandpass filters to select the emitted light, multiplexed imaging can be realized by changing the wavelength of the excitation light without the need for additional changes to the imaging interface [18]. Despite its advantages, optical microscopy in the tissue is constrained by the limited penetration depth of light delivered from external optical sources even with lower scattering and absorption of near-infra-red (NIR) wavelengths [19]. Therefore, to capture the immune response and tumor state changes in the tissue, proximity of the optical source to the imager and the target is necessary. Several previous works have developed implantable image sensors for high-resolution fluorescence microscopy inside the tissue for various biomedical applications including optogenetics, optical functional imaging, biosensing and neural activity recording but they haven’t demonstrated wireless power transfer for an untethered implantable sensor. These systems rely on external wiring for power and data transfer [20]–[23] or they are supplied by batteries [25] and hence are not practical for chronic real-time monitoring. A wireless thermoacoustic imager has been introduced [26] but it does not provide the sensitivity and specificity needed to track the immune response. Fiber optic probes have shown *in vivo* imaging but their utility for continuous monitoring is limited by the invasiveness of the process [27].

To address these challenges, a device containing a fluorescence image sensor capable of local illumination while relying on wireless power and data transfer is needed. Here, as a first step towards the envisioned system in Fig. 1(a), we demonstrate design and implementation of a proof-of-concept platform with the following key capabilities: (1) Wireless power and data transfer via ultrasound (US) to supply the image sensor and transmit data using a single US transceiver. (2) Chip-scale fluorescence microscopy to detect 100s of cells for capturing the immune response and cancer state changes, (3) Supply of high instantaneous power to a micro laser diode (μLD) with limited available power harvested from US to provide local illumination for the image sensor. While this work aims to demonstrate the feasibility of this platform, future work is required to prepare the device for the intended setup including biocompatible packaging and assembly of the μLD next to the imager [29]–[31] or use of light-guided plates [32]. An analysis of the differences between the current platform in Fig. 1(b) and the intended setup in Fig. 1(a) is included in the supplementary. An US link, as opposed to inductive coupling, optical or RF-based links [33], is chosen due to low tissue attenuation (0.5-1 dB/cm/MHz [34]) and higher Food and Drug Administration (FDA) regulatory limit for power flux density (720 mW/cm^2^ [35]), allowing for efficient wireless power transfer deep within the body. A single piezoceramic is controlled by the sensor for wireless power transfer and data communication up to a depth of 2 cm. The device powers up using wireless energy from the US link, and following commands encoded in the US transmissions proceeds to illumination using the μLD while simultaneously capturing an image from fluorescently labeled targets. To eliminate bulky optical lenses, a microfabricated angle-selective collimator is utilized to restrict the angle of incident light resulting in sharper images with higher resolution building on our previous work [36]. Next, it serially converts each pixel to a digital value and transmits data back to an external transducer via US backscattering. As a demonstration of functionality, in the wired mode, the imager array can capture *ex vivo* images of CD8 T-cell profiles consistent with images taken with a high-resolution fluorescence microscope. The overall performance of the systems in wireless mode is demonstrated by resolving features on a USAF resolution test target and transmitting the data from a depth of 2 cm.

**Fig. 1.**
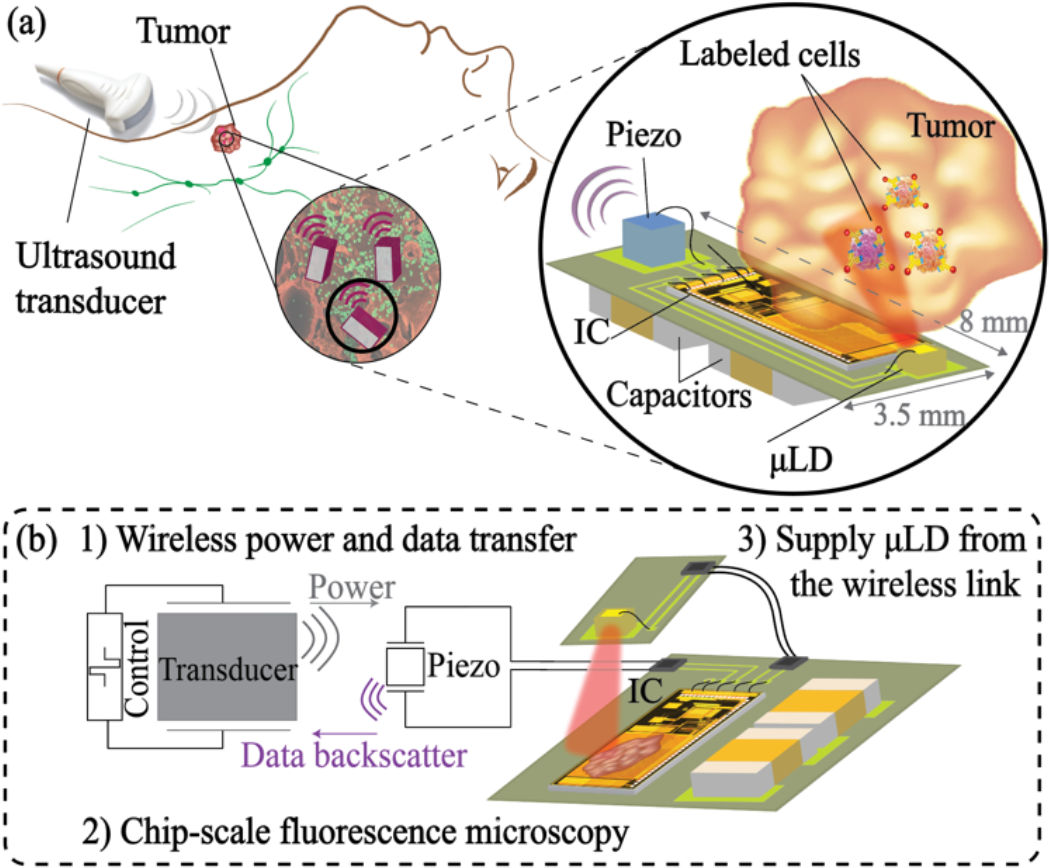
(a) Conceptual system diagram of the envisioned wireless imaging system in the tumor microenvironment with estimated sizing modified from [28]. (b) Major challenges towards (a) demonstrated in the current platform.

The paper further explains and expands the work presented in [28] and it is organized into seven sections. Section II outlines the challenges and requirements in wireless fluorescence microscopy for tracking the response to immunotherapy. Details of the wireless interface and strategies to reduce the size of the device are highlighted in section III. Section IV describes the design and operation of the subblocks including the image sensor and wireless power transfer with the backscattering protocol. Section V details *ex vivo* characterization of the imaging front-end in capturing cell clusters in the wired mode. We present experimental results validating the wireless system’s overall performance in section VI. Finally, the conclusion and comparison with the state-of-the-art are provided in section VII.

## II. Wireless Fluorescence Microscopy

To design a wireless system for fluorescence microscopy, as illustrated in Fig. 1, system-level integration of the IC and the optical source (as light cannot penetrate from an external source deep into the tissue) is essential. The miniaturized wireless system in Fig. 1 consists of 1) a 36×40-pixel lensless wireless CMOS imaging ASIC, 2) a sub-mm-sized laser diode, 3) a single 1.5×1.5×1.5 mm^3^ piezoceramic (Lead Zirconate Titanate, PZT) transceiver for wireless power harvesting and data transmission and 4) off-chip capacitors for energy storage. Quantifying the fluorescence signal from microscopic cell foci in immunotherapy is the key to determining the specifications of the device and the requisite optical power from the light source. This section describes the approach to quantify the fluorescence signal using biological samples and outlines the requirements for the optical source.

### A. Fluorescence Microscopy - Establishing Parameters for Quantitative Imaging

As shown in Fig. 2(a), to perform fluorescence microscopy, the targeted cells are labeled with fluorophore-labeled antibodies prior to imaging. This can be accomplished *in vivo* through an IV administration of the antibody-fluorophore conjugate targeting specific cell populations of interest [33]. The fluorophore determines the wavelength for imaging, and the antibody specificity ensures labeling the target cell type. Once bound to the cells, the fluorophores excited by the excitation light (shown in blue) emit photons with a longer wavelength (shown in red) after a small (<50 nm) Stokes shift as demonstrated in Fig. 2(b).

**Fig. 2.**
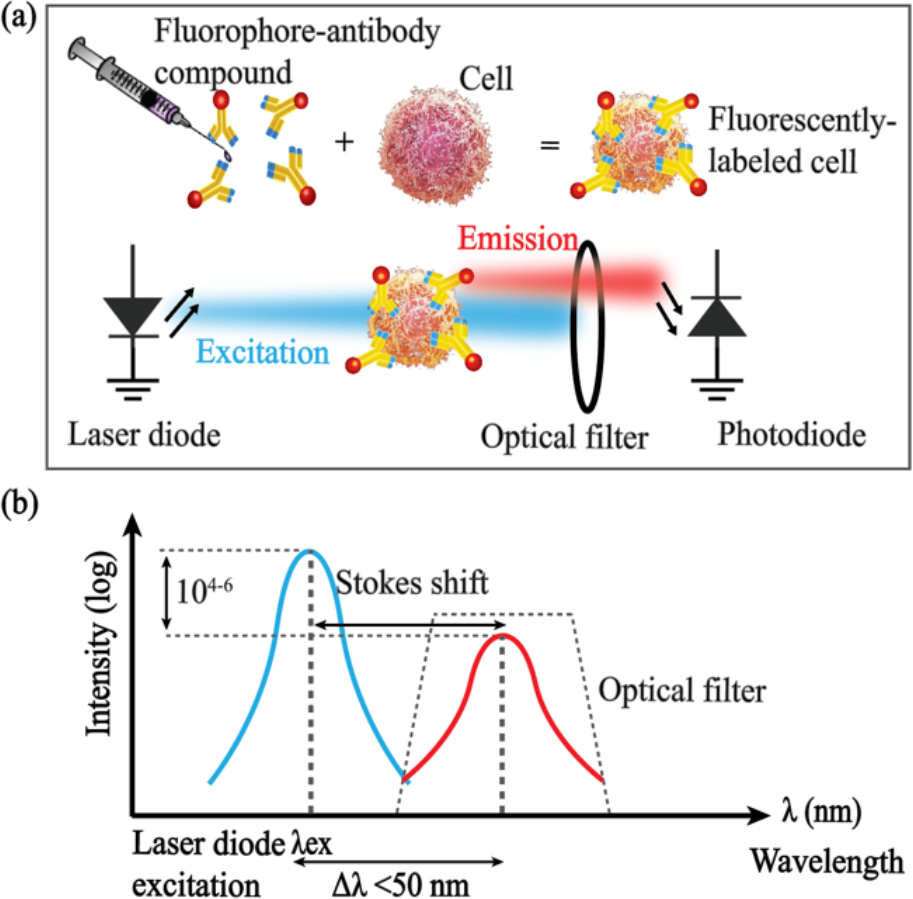
Fluorescence microscopy for fluorophore-labeled targets using a laser diode, an optical filter and a photodiode (from [28]).

The inefficiency of converting the excitation light to fluorescence [37] requires excitation intensities up to 6 orders of magnitude higher than the emission and a longpass or bandpass optical (wavelength) filter to attenuate the excitation light to prevent saturating the photodiodes in the sensor. Commonly used organic fluorescent dyes in biomedical studies (Cy5, IRDye800CW, etc.) feature Stokes shifts of only 20-30 nm [38], [39]. This small shift in the spectrum requires high-performance optical filters with high out-of-band attenuation (OD≥6) to eliminate bleed-through from the higher-intensity excitation (∼50 mW/cm^2^) to the emission band as the scattered excitation light contributes to the background signal.

### B. Fluorescence Signal Quantification

The fluorescence signal intensity from fluorophores bound to cell foci can be estimated given the excitation power density of the optical source, the number of fluorophores per cell, and the optical properties of the fluorophore (quantum yield and absorption cross-section).

The fluorescence signal from a cluster of cells is given by

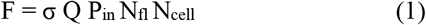

where σ is the absorption cross-section of the fluorophores in cm^2^, Q is the fluorescence quantum yield, P_in_ is the incident optical intensity in mW/cm^2^, N_fl_ refers to the number of fluorophores attached to the target cell and N_cell_ is the total number of cells in the cluster. Typical antibodies demonstrate an average binding of 10^6^ per cell [40], with 1-5 fluorophores per antibody, resulting in ∼10^6^ fluorophores per cell [41], a quantum yield of 20%, and an average absorption cross-section on the order of 10^−16^ cm^2^ [42], [43]. Characterizing the range of the input signal from the desired cell foci given the responsivity of the detector determines the power requirements of the optical source. For example, to achieve SNR>10 dB in detecting the fluorescence signal from a cluster of 500 cells, considered as a point source of light, 0.9 mm away from the sensor with a typical exposure time of 32ms using the pixel designed in [36], a high >50 mW/cm^2^ optical excitation intensity incident on the sample is required. The optical excitation intensity is more than 2 orders of magnitude lower than the maximum exposure limit imposed by the American National Standard for Safe Use of Lasers requirements as discussed in the supplementary section.

Of critical interest to assess therapy response is the *change* in both the number and position of cell. To estimate the change in the number of cells needed to evaluate the immune system’s response, we analyzed fluorescence images from an experiment in which immunotherapy is delivered to mice implanted with tumors. Fig. 3 shows images taken with a fluorescence microscopy scanner (Axio Scan.Z1, Zeiss) from sectioned lymph nodes of an untreated and treated mouse with immunotherapy, each harboring an implanted tumor (MC38 cell line). For each sample, CD8 T-cells, a principal component of the immune system’s response to cancer are stained (shown in pink) and demonstrated overlaid with the cell nuclei of the entire sample (DAPI stain shown in blue). Current studies in immunotherapy also support increasing populations of CD8 T-cells shown in the fluorescence images of the tumor microenvironment (TME) as an indicator of the immune response [44], [45]. Comparing the cross-sectional samples in Fig. 3 shows an increase of ∼500-1000 in the number of CD8 T-cells proliferating the lymph node of a treated mouse which is consistent with the estimated value for N_cell_ in (1).

**Fig. 3.**
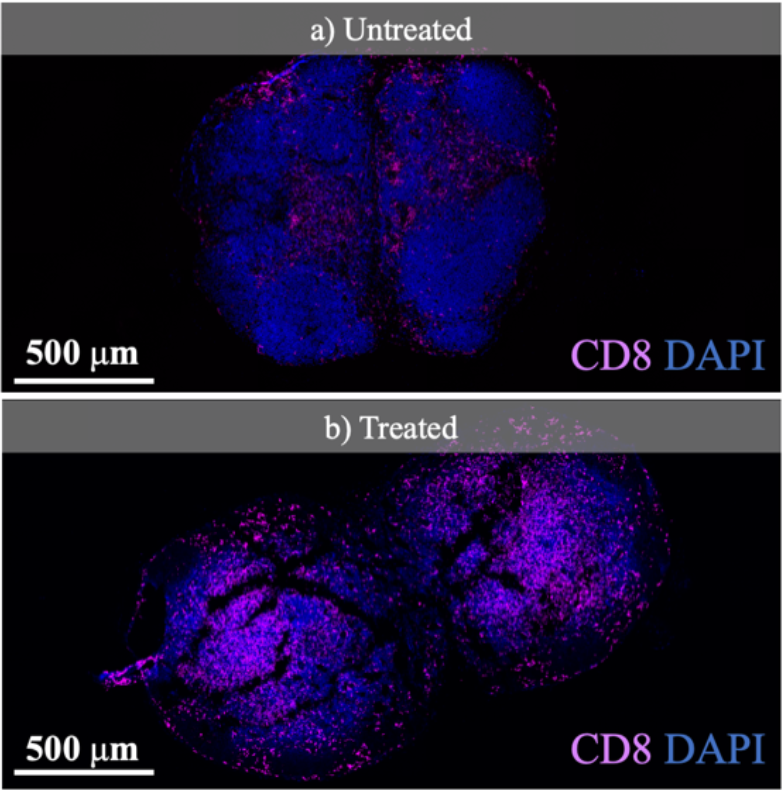
Fluorescence images of CD8 T-cells in the lymph nodes of an (a) untreated and (b) treated mouse with immunotherapy.

### C. Optical Power Requirements

In addition to the emitted photons, the scattered excitation light from the background contributes to the photodiode signal. Given the typical 20-30 nm Stokes shift of the common organic fluorophores and the high excitation intensity (∼50 mW/cm^2^), even using a filter with out-of-band transmission of up to 6 OD can result in bleed-through of the excitation light and saturation of the pixels. To obviate the need for excitation filters, a laser diode is chosen over an LED. In order to achieve deeper penetration of the excitation and lower tissue autofluorescence [19], [46], a laser diode with a peak wavelength of 635 nm is chosen. The electro-optical PIV characterization of the edge emitter μLD (0.3×0.25×0.1 mm^3^, CHIP-635-P5, Roithner Laser Technik) using a power meter (PM100D, Thorlabs) is shown in Fig. 4. The nominal forward voltage and current of 2.1 V and 37 mA, respectively, result in a total measured optical power of 3.4 mW and an electrical to optical efficiency of 4.4%, necessitating significant power delivery to the sensor for fluorescence imaging. Details of supplying the μLD while imaging the samples are included in section III.

**Fig. 4.**
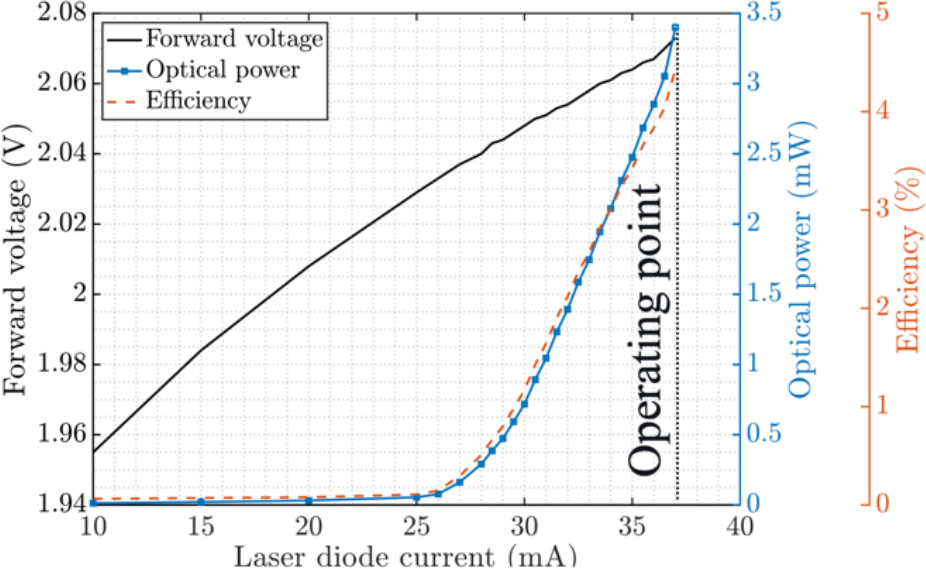
PIV characteristic of the micro laser diode (µLD) (0.25×0.3×0.1 mm^3^, CHIP-635-P5, Roithner LaserTechnik) from [28].

## III. Wireless Fluorescence Microscope

Given that the available power from US is smaller than the required 78 mW instantaneous power of the μLD, imaging is performed in a duty-cycled fashion. Thus, the *Charge-Up* state (when the required energy for taking an image is stored) is separated from the *Imaging* state (when the stored energy is used to illuminate the sample and capture the image) in the time domain. This section describes the system-level requirements of the imager array and the acoustic interface for wireless chip-scale fluorescence microscopy.

### A. Imager Array for In Vivo Microscopy

The design of the imager array is based on our previous work in [36]. Each pixel incorporates a 44×44 µm^2^ photodiode and has a pitch of 55 µm. The pixels are fabricated in a 0.18 µm 1.8/5/32 V TSMC CMOS process with a dark current density of 14 aA/µm^2^ and a responsivity equal to 0.09 A/W at 635 nm. To obviate the need for bulky optical lenses, on-chip microfabricated structures based on angle-selective gratings (ASGs) with FWHM of 36° are utilized to restrict the angle of incident light resulting in images with higher spatial resolution [36]. The use of ASG, along with in-pixel electronics, yields an effective fill factor of 28%. Imaging is performed using a global shutter as the LD only illuminates the sample for a limited time, which is by far the primary power-consuming operation of the imager, and therefore all pixels must image during this limited time window. This demands that each pixel be able to amplify, sample and hold its data until it is read out via a single channel US-based uplink which will be discussed in section IV. Each pixel consists of a capacitive trans-impedance amplifier (CTIA) with a custom-designed integration capacitor C_int_=11 fF. The photodiode signal of each pixel can be computed from V_signal_ = i_pd_T_int_/C_int_, where i_pd_ is the photodiode current and T_int_ is the integration (exposure) time [36].

The pixel size (*W*_*pixel*_) and the integration time (*T*_*int*_) are chosen to maximize sensitivity to the dynamics of small cell foci (a few 100 cells) to evaluate the immune response. To capture cell movements in real-time, the pixel must be small enough to track the displacement of cells within consecutive frames. The minimum interval between frames is constrained by the charging time of the storage capacitor as it is used to supply the energy for the μLD during the *Imaging* state. Therefore, given a constant power consumption of the μLD, the minimum frame time is a function of the time it is switched on, T_int_. For each T_int,_ the pixel dimension must be consistent with the typical displacement of cells with an average velocity of 10-20 μm/min ([47],[48]) between each frame. Pixels with dimensions much larger than this displacement may miss the changes in the cell proliferation profile. Conversely, designing an imager array with the same imaging area using smaller pixels results in unnecessarily oversampling the scene. As discussed in [40] there is a fundamental tradeoff for *W*_*pixel*_ between maximizing the signal and maintaining spatial resolution. The received fluorescence signal is proportional to the active area of the photodiodes until the field of view of a pixel matches the size of the foci being imaged. It should be noted that the goal is to track changes in cellular distribution (in response to therapy), and not to obtain intracellular or single-cell imaging. Thus, too small of a pixel will capture noise with only a minimal detected signal which results in a low SNR. The typical ∼10 μm dimension of each cell introduces a lower bound for the pixel size. Given the size of the pixel’s peripheral circuitry needed to ensure low noise in-pixel amplification and sample and hold, shrinking *W*_*pixel*_ lowers both the fill factor and the sensitivity to light. Subtending the same field of view inside the TME with smaller pixels results in larger arrays with higher power consumption and longer readout times. The spatial resolution depends on both the resolution of the angle-selective structures as well as the pixel dimension, and therefore lowering the pixel size significantly beyond the optical resolution will not result in further improvements in resolution. Conversely, larger pixels that reduce spatial resolution collect more dark current in addition to the photodiode signal, both as a linear function of the area. Given that dark current in this technology is the dominant factor restricting the dynamic range of the pixel, larger pixel area results in higher photodiode shot noise limiting SNR and thereby the minimum detectable signal.

To quantify the trade-offs listed above a figure of merit (FoM) is defined to incorporate the specifications of both the imager array and the wireless system. This proposed metric consists of1) SNR to ensure sufficient image quality while capturing multicellular dynamics, 2) spatial resolution to enable resolving small feature sizes, and 3) overall form factor to facilitate eventual implantation. Given the importance of image quality (SNR, resolution), and, secondarily the need to miniaturize the form factor of the implant, we propose an FoM given by

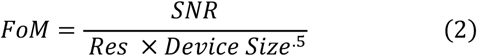

Optimization is performed on *W*_*pixel*_ and *T*_*int*_. The power of each term in (2) is chosen to balance dependency on the order of the independent variables. The size of the device is dominated by the storage capacitor which, given a constant LD power, is proportional to the integration time *T*_*int*_. Resolution is proportional to the pixel dimension and SNR is defined as

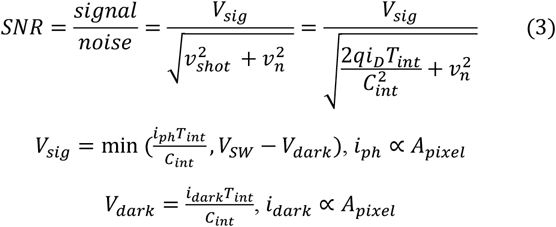

where *v*_*n*_ refers to the overall noise contribution of the in-pixel circuitry excluding the shot noise of the photodiode represented as *v*_*shot*_. A detailed quantification of pixel noise is presented in [36]. The photodiode current signal is *i*_*ph*_, the dark current is *i*_*dark*_, *q* is the charge of an electron and *i*_*D*_is the total current through the photodiode including both the signal and dark current (*i*_*D*_ = *i*_*ph*_ + *i*_*dark*_). Both the photodiode and dark current are proportional to the area of the pixel 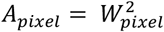 for square-shaped pixels with a width of *W*_*pixel*_. *V*_*SW*_ is the maximum voltage swing at the output of the pixel. The maximum voltage generated from the photodiode signal is constrained by the contribution of the dark current in the pixel output voltage (*V*_*dark*_). This introduces an upper bound for the maximum signal that can be detected by each pixel. The spatial resolution of the imager is defined by the dimension of the pixel and the optical angle selective structures. Assuming that the optical structures provide sufficient resolution, the overall resolution is limited by *W*_*pixel*_. Fig. 5 demonstrates the FoM defined in (2) and its contours for combinations of *W*_*pixel*_ and *T*_*int*_. The blurred region corresponds to the design space resulting in SNR values lower than 10 dB which lack adequate image quality for accurate detection of the desired cell clusters. The dashed line represents the lower bound of the pixel size needed to capture cell displacements in consecutive frames. This is computed given average cell velocities (10-20 μm/min) and the minimum achievable frame time for each *T*_*int*_. Oversampling the scene with an imager made with smaller pixels than the lower bound increases the number of pixels, and thereby the power consumption and data transmission period. To maintain adequate resolution in detecting multicellular-level dynamics in the TME, an upper bound on the pixel size is defined as highlighted in Fig. 5(b).

**Fig. 5.**
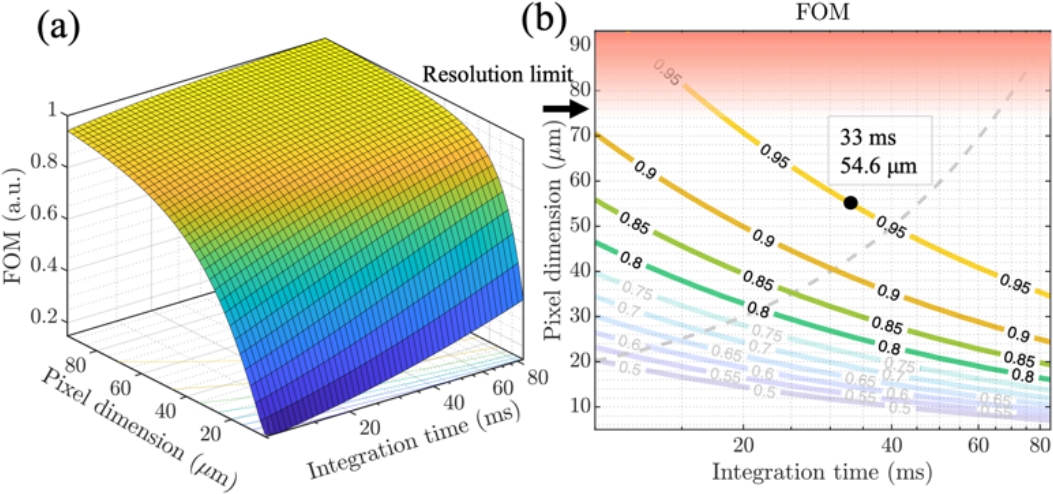
FoM optimization based on pixel size (W_pixel_) and integration time (T_int_). (a) Normalized FoM for the overall design space. (b) FoM contours with 1) blurred regions corresponding to SNR<10 dB, 2) dashed line representing the sufficient lower bound W_pixel_ for each T_int_ according to cell dynamics and 3) highlighted region for the upper bound on W_pixel_ to ensure sufficient resolution.

According to the FoM optimization analysis, an imaging array of 36×40 pixels with a 55 µm pitch is chosen to fit a 44×44 µm^2^ photodiode, an in-pixel amplifier, and sample and hold circuits for each pixel. To visualize multicellular clusters of a few 100 cells illuminated with the μLD using these pixels, an integration time equal to 32 ms is chosen to optimize the FoM within 95% of its maximum. With T_int_ programmability introduced in section IV, our proposed system can maintain a high FoM tailored to capture different cell profiles and fluorescence signal intensities in the TME.

### B. Wireless Link

The operation of the chip is divided into multiple states. For each frame, *Imaging* only occurs when there is enough charge stored in the off-chip storage capacitor, C_store_, to supply the laser diode during the entire integration time. Various blocks including the μLD driver and the pixel array are power gated to only turn on during their operating states to minimize power consumption which is explained in section IV. To derive requirements for C_store_ and the wireless link, the simplified model of the chip in Fig. 6 can be used. The laser driver is modeled with a current source turned on only during the *Imaging* state (T_int_). During *Imaging*, the current of the laser diode, *I*_*LD*_, dominates the current consumption of the rest of the chip modeled with a single load resistance *R*_*L*_. Given the availability of 5 V devices in the CMOS process, the maximum rectified harvested voltage (V_rect_) is set to 5 V. The maximum voltage drop on the storage capacitor during *Imaging* is limited to 1.5 V as a lower bound of 3.5 V is needed to ensure a minimum supply voltage for chip operation after the *Imaging* state. To attain the 37 mA nominal current for the μLD (*I*_*LD*_) required for imaging the immune system components during a 32 ms integration time with a maximum voltage drop ΔV = 1.5 V, 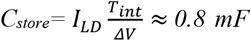 is chosen. The value of C_store_ can be decreased in the future by utilizing more efficient optical sources or by allowing a larger voltage drop on V_rect_.

**Fig. 6.**
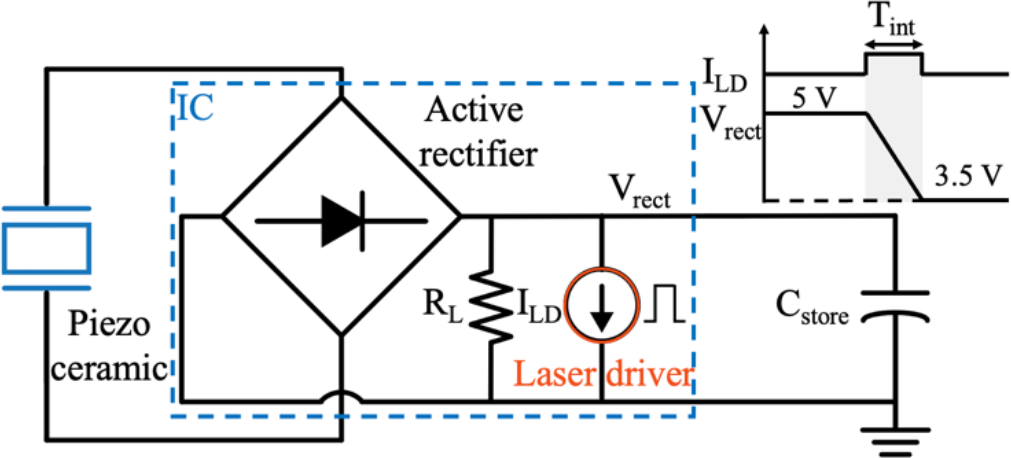
Simplified block diagram during the *Imaging* state. Maximum 1.5 V linear voltage drop on V_rect_ while supplying the laser driver from C_store_.

Another approach to lower C_store_ without compromising the performance of the device is to shorten T_int_ and at the same time take multiple averages to maintain the initial SNR while keeping I_*LD*_ and therefore the optical power of the μLD constant. For a CTIA-based pixel, the electrical noise component consists of the shot noise of the photodiode, the thermal and flicker noise of the transistors in the pixel amplifier, and the thermal noise of the sample and hold switches. Fixed pattern noise from dark current and pixel variation can be eliminated by subtraction of the dark imager or calibration of the pixels, hence they are not included in the analysis. As shown in (3), lowering T_int_ results in SNR degradation. To maintain SNR after decreasing T_int_, averaging can be applied to reduce uncorrelated noise by a factor of √n where n is the total number of averages. For the overall measurement time to stay constant, the number of averages can be scaled by the same ratio of downsizing T_int_. SNR of the image with an integration time of T_int_/n after taking n averages is given by

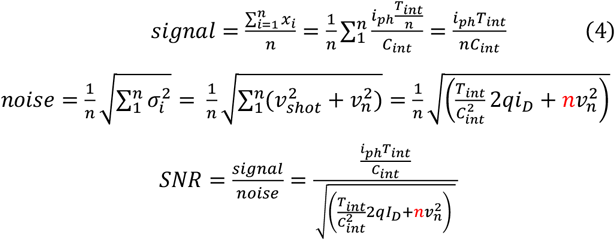

where *x*_*i*_ refers to the pixel value from each measurement assuming that the voltage swing is not saturated by the dark current. For a shot noise-limited design (*v*_*n*_<<*v*_*shot*_), averaging the images by the same factor of lowering T_int_ preserves SNR, while needing less energy for μLD for each image– reducing the size of C_store_, which drives the overall device from factor. In a shot noise-limited region, SNR is not affected by lowering T_int_ as long as the images are averaged accordingly. However, further reduction of T_int_ results in SNR degradation as the pixel amplifier noise becomes the dominant noise component (*v*_*n*_ >>*v*_*shot*_). Since the current pixel architecture is shot noise limited, the size of the storage capacitor can be reduced by a factor of 2-3 before SNR decreases by more than 1 dB.

As will be shown in section IV, device *Charge-up* dominates the frame time and is a linear function of C_store_. While this approach can reduce C_store_ and therefore the frame time by n for a single image, the overall measurement time for acquiring n frames will roughly stay constant. Consequently, the energy requirements from the US source are approximately constant. As lowering T_int_ continues and amplifier noise becomes the dominant noise component, maintaining SNR requires more averages (> *n*) resulting in an increase in the overall measurement time and energy requirements of the device.

To capture the movements of cells in the TME (∼10-20 μm/min) with the optimum value for the pixel size (55 μm) the frame time of the sensor must be less than a few minutes. A piezoceramic with a size of 1.5×1.5×1.5 mm^3^ is chosen to charge C_store_ with the energy from the acoustic waves. Fig. 7 shows the impedance and harvested open circuit voltage of the piezoceramic vs. frequency inside canola oil at a depth of 2 cm, with an acoustic attenuation (∼0.25 dB/cm/MHz) mimicking the acoustic properties of tissue. The frequency dependency of the harvested voltage is shown for the piezoceramic both without loading and loaded with the equivalent model of the chip. The open circuit voltage is measured using a focused acoustic transducer (V303-SU-F0.80IN-PTF, Olympus). Since V_rect_ cannot drop below 3.5 V to maintain chip operation, we maximize the peak harvested voltage on V_rect_ to allow for minimizing the size of C_store_ for the same acoustic flux. To maximize V_rect_ according to the resonance spectrum of the piezoceramic and the US transducer, the frequency of operation is tuned in between the series and parallel resonance frequencies of the piezoceramic at 960 kHz. In order to obtain V_rect_ = 5 V, the acoustic power flux density was increased to 905 mW/cm^2^ which exceeds the FDA-approved limits by 26%. In the future, this can be mitigated by lowering the required harvested voltage and using a voltage multiplier to reach the final 5 V.

**Fig. 7.**
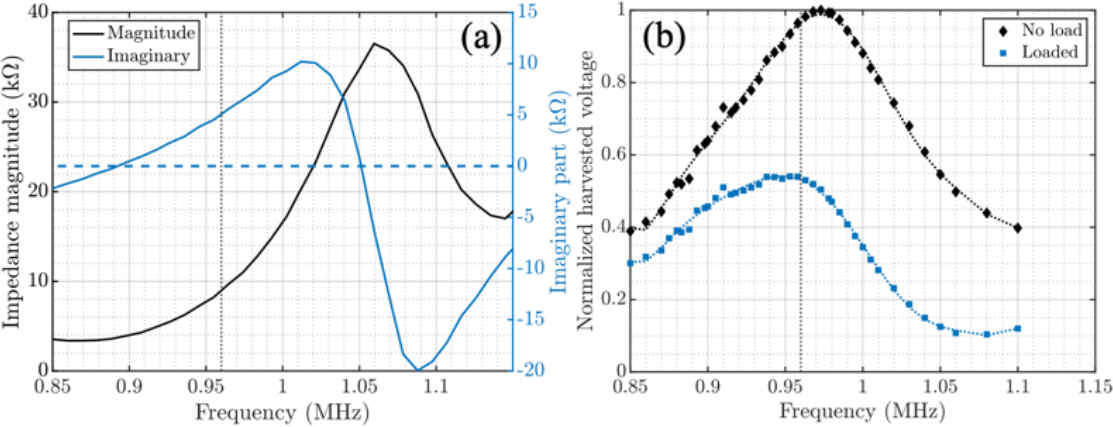
Characterization of the piezoceramic: Frequency spectrum of (a) magnitude and imaginary part of the impedance and (b) normalized harvested voltage of the piezoceramic with no load vs. being loaded with the equivalent of the chip’s input impedance from [28].

## IV. System Design and Implementation

Fig. 8(a) shows the block diagram of the ASIC including 4 main functional blocks: (1) power management unit (PMU), (2) imaging front-end, (3) laser driver, and (4) finite state machine (FSM). The PMU incorporates an active rectifier and several low-dropout voltage regulators (LDOs) to supply various subblocks. The imaging front-end consists of the pixel array shown in Fig. 8(b) with the architecture of the pixel and the sample and hold in Fig. 8(c) and (d) as discussed in [36]. The imager array is followed by the readout circuitry, buffers, and a differential 8-bit SAR ADC for reading out and digitizing the analog pixel values as illustrated in Fig. 8(a). The laser driver supplies a constant current to the laser diode from the charge stored in C_store_. The FSM controls the timing and operation of the chip and synchronizes it with the external US transducer.

**Fig. 8.**
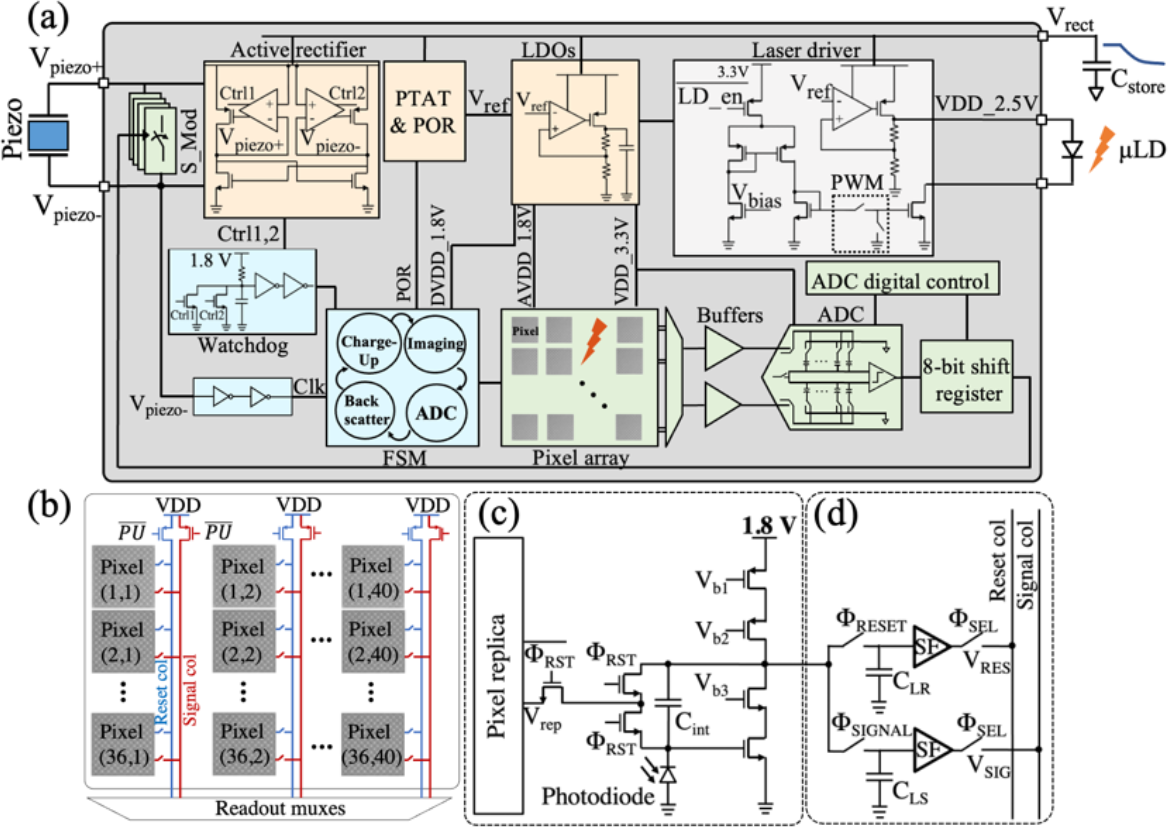
(a) Block diagram of the IC including power management, imaging front end, laser driver, and FSM. The IC is connected to the piezoceramic, external C_store_, and μLD [28]. (b) The architecture of the imager array with the shared reset and signal within a column and readout (c) Schematic of a pixel including the photodiode, the pixel CTIA and the replica circuit as discussed in [36] and (d) In-pixel sample and hold of V_RES_ and V_SIG_ from correlated double sampling.

The micrograph of the ASIC appears in Fig. 9. The chip measures 2.4 mm by 4.7 mm with the pixel array taking up 41% of the overall area. The design and operation of each block are described in detail in the following:

**Fig. 9.**
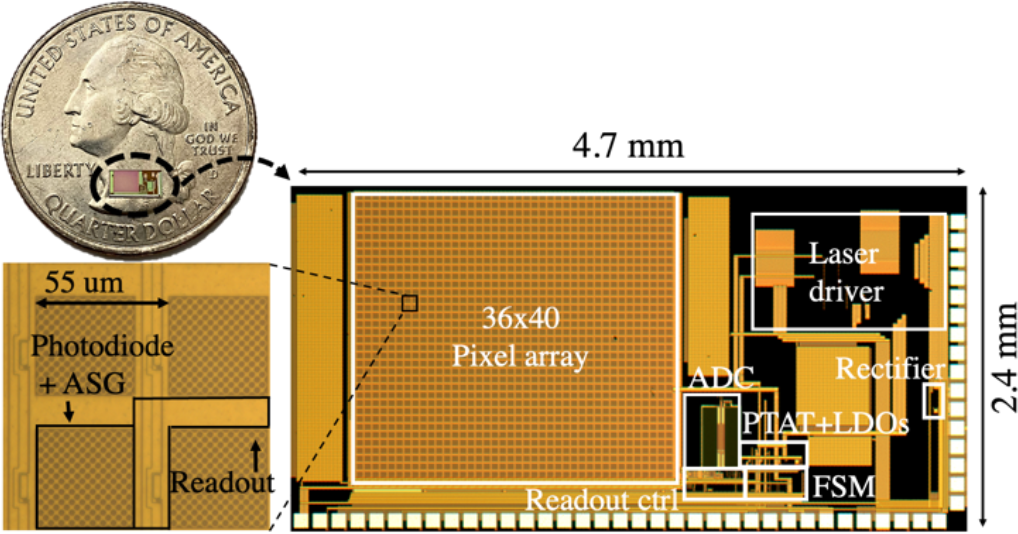
Chip microphotograph showing the imager array, readout digital control, ADC, rectifier, PTAT, voltage regulators, FSM and laser driver. The chip is 2.4 mm by 4.7 mm and the pixel array measures 2 mm by 2.2 mm. Close-up view of the pixel with a 55 μm pitch including a 44 μm by 44 μm photodiode area covered with ASGs and the readout circuitry (from [28]).

### A. PMU and Digital Control

As shown in the timing diagram in Fig. 10, the operation of *the* chip is divided into 4 states: *Charge-Up, Imaging, ADC Operation*, and *Backscatter Modulation*. To eliminate the complexity of data downlink and ensure that on-chip state transitions are synchronized with the transducer, the transmitted US carrier is modulated with a pulse sequence. The different pulse widths of the US signal for each state of operation are programmed with an FPGA which controls the output of the US transducer shown in Fig. 10 (V_piezo+_). A watchdog control signal demodulates the incoming US waveform’s envelope to navigate the state transitions of the FSM.

**Fig. 10.**
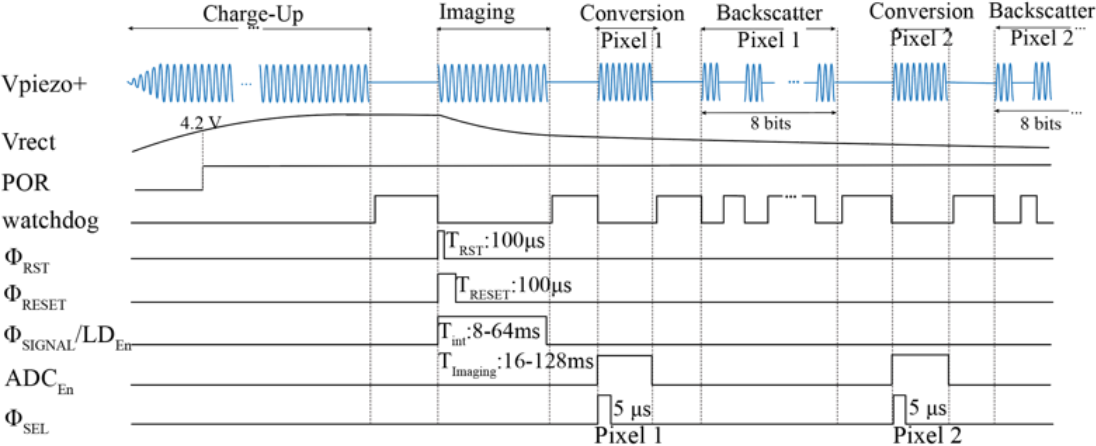
Timing diagram and state transitions of the system based on the pattern of the ultrasound waveform received by the piezoceramic (V_piezo+_). The timing control signals for each pixel’s operation (Φ_RST_, Φ_RESET_, Φ_SIGNAL_) are shown. LD_En_ and ADC_En_ refer to the control signals to turn on the laser driver and ADC, respectively. Φ_SEL_ indicates the sampling phase of the ADC when the pixel’s dual outputs are being read (from [28].)

The active rectifier converts the US signal to a DC voltage (V_rect_) while charging C_store_ up to 5 V. To initialize the chip and reset the FSM, an on-chip power-on reset (POR) signal is triggered as V_rect_ reaches 4.2 V to guarantee that the LDO voltages are established. Various on-chip LDOs (1 V, 1.8 V, 2.1 V, 2.5 V, 3.3 V) with a total current consumption of 8.2 μA regulate the supply voltage for the analog front-end, the laser driver, and the FSM. Despite the droop in V_rect_ during the *Imaging* state, the LDOs are designed to operate with V_rect_ as low as 3.5 V to ensure the functionality of the device after *Imaging* for *ADC Operation and Backscatter Modulation*. A CLK signal with a frequency of 960 kHz is extracted directly from the acoustic carrier.

Power-intensive blocks including the laser diode driver, the pixel array, the ADC, and the buffers preceding the ADC are turned off during *Charge-Up* to prevent disrupting and extending the chip’s power-up. The enable signals for the laser diode driver (LD_En_), ADC (ADC_En_), and ADC buffers (Φ_SEL_) are shown in Fig. 10. Followed by the *Imaging* state, the first rising edge of the watchdog signal is indicative of the end of the *Charge-up* period. The exact duration of the *Charge-Up* state can be empirically determined by characterizing the rise time of V_rect_ to reach its final value (5 V) for a given C_store_. During *Imaging*, the sample is illuminated by the sensor-powered laser diode and after the image is captured, the pixels are read out, digitized, and backscattered sequentially. Each pixel’s voltage is wirelessly transmitted by modulating the impedance of the same piezoceramic used for power transfer. Data transmission continues until the watchdog timer counts the entire 1440 pixels based on the transitions of the watchdog signal. The data transfer protocol is discussed in part C.

### B. Imaging and Laser Driver Operation

During *Imaging*, the photodiodes convert incoming photons from the fluorescently labeled cells into a photocurrent, which is integrated into the feedback capacitor of the pixel CTIAs, C_int_ as shown in Fig. 8(c). The output voltage is sampled twice, once at the beginning (V_RES_), and again at the end of the integration time (V_SIG_) generating reference and signal values respectively, which are subtracted from each other to provide the net signal. This correlated double sampling (CDS) approach in Fig. 8(d) suppresses offset and low-frequency noise of the pixel. The pixel array is turned on only during the *Imaging* state when it consumes a total current of 145 μA. A detailed design of the pixels is presented in our previous work [36]. The laser driver schematic is shown in Fig. 8(a) [49]. To prevent the LD from overheating, the driver supplies the laser diode with 50% duty-cycled 50 kHz current pulses as opposed to a continuous current. Therefore, the integration time is effectively half of the duration of the *Imaging* state (T_int_=32 ms for a 64 ms *Imaging* state). A PWM controller sets the frequency and duty cycle of the pulses based on the main CLK frequency. The output of the PWM block drives a complementary set of switches to control the current of the laser driver. The supply voltage of the laser driver is regulated to 2.5 V to comply with the maximum voltage allowed for the laser diode. A small off-chip resistor in series with the laser diode can adjust the voltage in case of variations. Based on the signal intensity and size of C_store_, 8 integration times ranging from 8 ms to 64 ms in steps of 8 ms can be configured into the chip at the package level.

### C. Data Conversion and Backscattering

Once the image is captured, both the laser driver and the pixel array are switched off and the FSM transitions lead the chip to *ADC Operation* and *Backscatter Modulation* states. At the beginning of each ADC state, the correct row is selected by digital row-driving circuitry. In each row, the reset and signal voltages of each pixel from CDS are read sequentially and sampled during a 5 μs sampling phase of the differential ADC. During this sampling phase referred to as Φ_*SEL*_ in Fig. 10, the ADC input buffers turn on and the readout circuitry selects the correct pixel from the imager array. The output of the ADC is serialized with an 8-bit shift register and is backscattered using pulsed-echo on-off keying (OOK) modulation to sustain a low bit error rate (BER). Backscatter modulation is realized by altering the electrical load resistance of the piezoceramic which affects the acoustic reflection coefficient of the incident acoustic signal [50]. The pulsed-echo modulation scheme is implemented to separate power and data transfer in the time domain while using a single piezoceramic for both.

The proposed backscatter modulation scheme is shown in Fig. 11. For a depth of 2 cm, the 8-bit packet of each pixel is divided into 4 sets of 2 bits fit within the 26.7 us roundtrip (=2ToF, time-of-flight) of the acoustic waves in oil. The US transducer interrogates the piezoceramic with the modulated waveform shown in Fig. 11. After each sequence of 2 bits, the transducer stops interrogating for 2ToF, to eliminate interference from the high voltage power waveform with the weaker backscattered signals. Once the signal reaches the piezo after a single ToF, it is modulated based on the acoustic reflection coefficient resulting from the impedance of the chip, R_Load_. At the series (fs) and parallel (fp) resonance frequencies, the normalized backscattered echo amplitude is proportional to R_Load_/(R_Load_+R_piezo,s_) and R_piezo,p_/(R_Load_+R_piezo,p_), respectively, where R_piezo,s_ and R_piezo,p_ are the equivalent resistances of the piezo at fs and fp. For the rest of the frequencies, the reflection coefficient can be computed given the piezoceramic properties and R_Load_ as quantified in [50]. A modulation switch, S_Mod in Fig. 8(a), is used to modulate R_Load_ and ultimately the echo amplitude for OOK modulation. A programmable switch with 4 impedance values (1,2,4,8 kΩ) sets the modulation depth based on the piezoceramic’s equivalent impedance at the operating frequency. Finally, after the second ToF, the backscattered signal appears on the transducer which is now in the receiving mode and will be demodulated and post-processed in MATLAB to reconstruct the image.

**Fig. 11.**
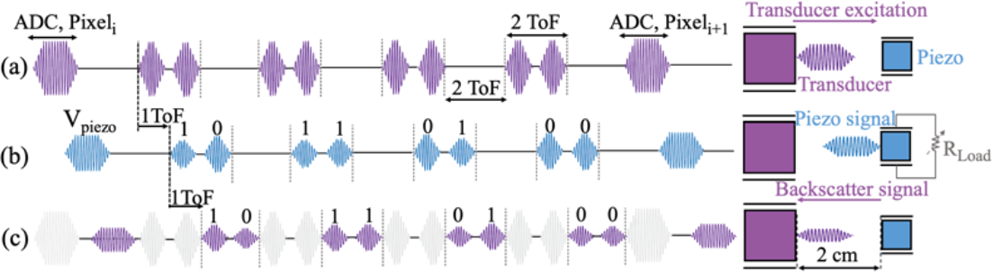
Backscatter modulation scheme (a) The US transducer interrogates the piezoceramic with a sequence including waveforms for the *ADC Operation* and *Backscatter Modulation* states (b) After 1 ToF, the piezoceramic receives the signal and modulates the pulses with R_Load_ according to each bit’s value (c) The backscattered signal is received by the transducer after a second ToF. The amplitude of the backscattered bits depends on the reflection coefficient of the acoustic waves as a function of R_Load_ and the frequency of operation.

For each pixel, the FSM alternates between *ADC* and *Backscattering* states until all the image data for a single frame is transferred. The conversion and backscattering for all pixels take 389 ms for an implantation depth of 2 cm. The received backscattered waveform is filtered by an FIR bandpass filter in MATLAB to improve signal quality. For measurement performed inside oil at V_rect_ = 5 V (when taking a dark current image and turning off the laser driver during *Imaging*) the modulation depth is 14.1% and data transmission is error-free for 11.52 kbits of data transmitted resulting in a bit error rate (BER) of better than 8.68×10^−5^. Lowering V_rect_ to the minimum of 3.5 V decreases the modulation depth to 7.4% increasing the BER to 3.47×10^−3^ for the transmitted data (11.52 kbits). The histograms of the 0 and 1 bits for each value of V_rect_ are shown in Fig. 12. Future work to add error correction codes to the on-chip transmitter [51] or further averaging the image can reduce the error in the final reconstructed image.

**Fig. 12.**
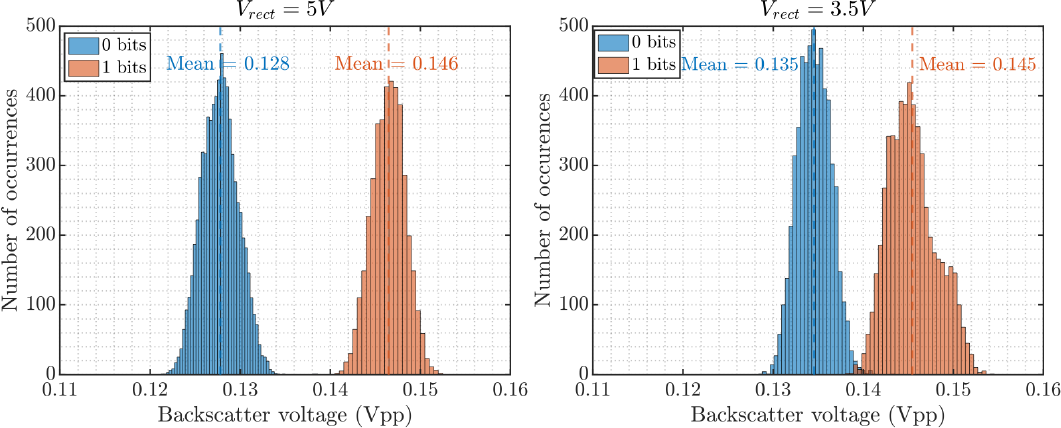
Modulation depth of 14.1% and BER better than 8.6×10^−5^ (error-free 11.52 kbits of data) for Vrect = 5 from [28] and modulation depth of 7.4% and BER of 3.47×10^−3^ for V_rect_ = 3.5 V.

Fig.13(a) shows state transitions of the chip after a 150 s *Charge-Up* state. A 64 ms *Imaging* state (T_int_=32 ms) and a portion of the *ADC* and *Backscatter Modulation* states are shown. After a linear 1 V drop during the *Imaging* state, due to the use of a larger 1.2 mF storage capacitor to lower the voltage drop, V_rect_ maintains a voltage higher than 3.5 V. In Fig. 13(b), the *ADC* and *Backscattering* states are shown for a single pixel with the modulated piezo signal corresponding to the bit values. The laser driver’s current is measured through the voltage across a 1.4 Ω resistor in series with the μLD. The laser driver current varies from 38.5 mA to 34.3 mA with a mean of 36.1 mA as V_rect_ drops from 5 V to 3.5 V. These currents correspond to optical powers ranging from 4.4 mW to 2.2 mW with a mean of 2.9 mW. The 11.5% current drop throughout the *Imaging* state stems from the drop in V_rect_ leading to a change in the PTAT output current which determines the laser driver current. This can be improved by using a larger storage capacitor. This results in the laser driver’s average electrical efficiency of 50%.

**Fig. 13.**
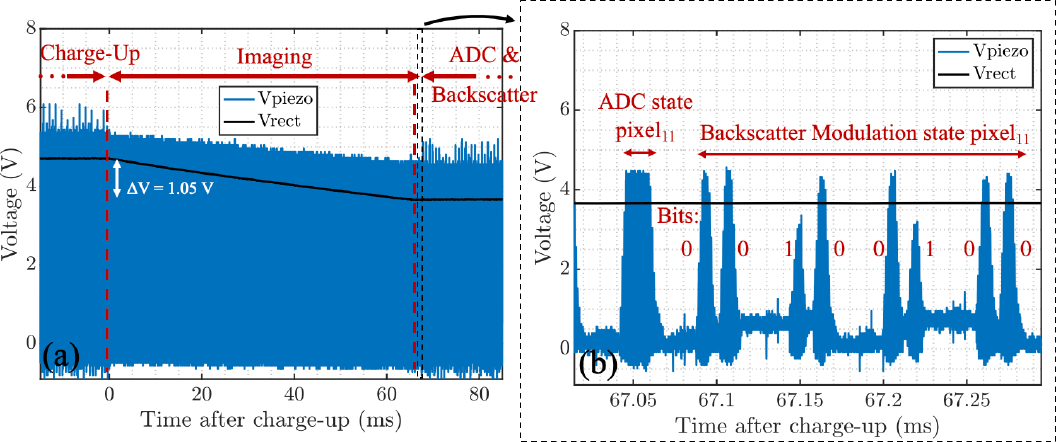
(a) State transition including the last part of the *Charge-Up, Imaging*, and initial part of the *ADC* and *Backscatter Modulation* states. V_rect_ drops linearly during the 64 ms *Imaging* state with an effective 32 ms integration time (T_int_). (b) Detailed transient waveform during the *ADC* and *Backscatter Modulation* states for each pixel from [28].

## V. Wired Mode Immune System *E**x* *V**ivo* Microscopy

The performance of the imaging front-end is tested by imaging *ex vivo* samples of a mouse model of cancer in an experiment monitoring the response to immunotherapy over 18 days.

### A. Experimental Setup

During the therapy, a group of 30 mice (strain: 006772) with a functional immune system (BL6 mice) against cancer (colorectal cancer, MC38 cell line) is selected. For each mouse, the tumor is implanted by injecting 5 × 10^5^ cells in 100 μL of MC38 cells in each flank. Once the tumors reach an appropriate size (∼ 5 mm), the mice are injected with 200 μg of immune checkpoint inhibitors, anti-PD1 and anti-CTLA4 [52],[53], two of the critical therapeutics that activate the immune system against cancer. The injections were repeated every 2-3 days, and 3 mice are collected at each serial time point (spanning 18 days with injection only happening during the first 12 days). The experiment is conducted under IACUC (Institutional Animal Care and Use Committee) protocol AN194778. At each time point, for each of the 3 mice, the draining lymph nodes from the tumor are harvested, fixed in formalin, embedded with paraffin and stained for CD8 T-cells with the Ventana Discovery Ultra automated slide stainer. To comply with the laser diode excitation wavelength, the samples are stained with the Cyanine5 (Cy5) dye-labeled antibodies targeted CD8 T-cells with absorption and emission peaks at 651 nm and 670 nm, respectively. As a proof of concept, CD8 T-cell populations in the lymph nodes of the untreated mice (day0, before injection, n=3) as controls and mice at the latest timepoint (day 18, n=3) are imaged with our proposed sensor and a fluorescence microscopy scanner (Axio Scan.Z1, Zeiss), to provide the ground truth.

The experiment setup is shown in Fig. 14(a). Illumination is provided with the 635 nm μLD controlled and powered by the chip which illuminates the sample from the top via transillumination. The bottom electrode of the μLD is mounted using conductive epoxy and the top electrode is wire bonded to the board. In future work, the μLD will be mounted on the same platform as the sensor. A 500-μm thick chip-size optical filter (ET FITC-Cy5, Chroma, EM: 675-755 nm) is epoxied on the chip using Sylgard 184 Silicone Elastomer mixed with a 1:10 ratio, degassed to remove any bubbles under a vacuum desiccator (SP Bel-Art) chamber, and then cured at 100°^C^ for 45 minutes. The filter demonstrates OD>6 at the excitation wavelength. More details on the optical front-end are included in our prior work [18]. The edges of the imager are covered with black epoxy (EP1046FG, Resinlab) (1:1 mixing ratio, cured at 65°^C^ for 30 minutes) to eliminate bleed-through from the sides and reflection from the wire bonds as shown in Fig. 14(b).

**Fig. 14.**
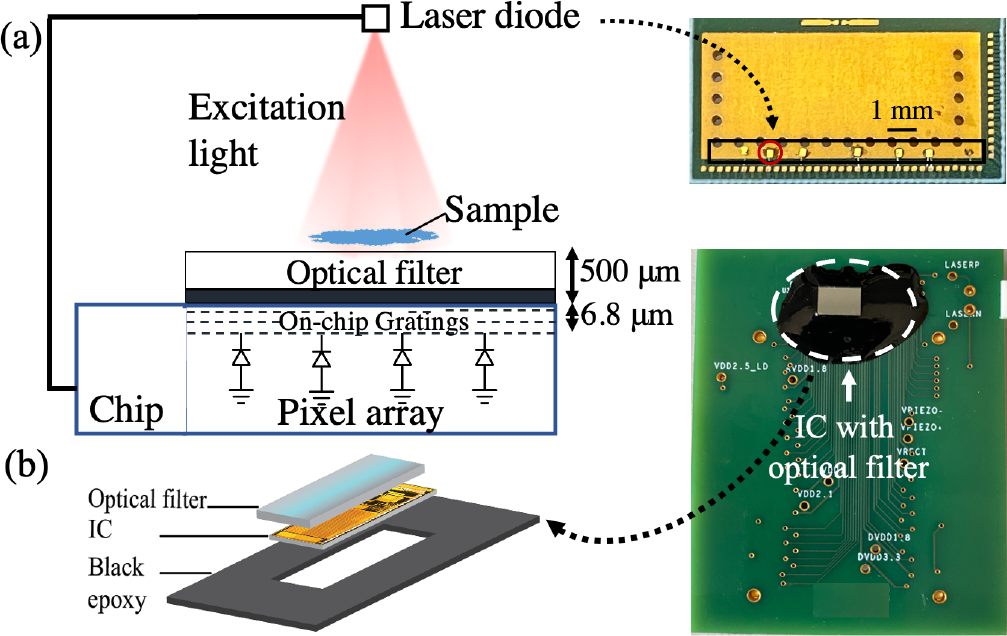
(a) Measurement setup for *ex vivo* microscopy in wired mode. The chip is powered up using a 250 kHz sinusoidal from the function generator instead of the 960 kHz piezo input. The on-chip laser driver controls the μLD which transilluminates the sample from the top. (b) Fabrication of the imager with the optical filter on the top and black epoxy on the sides to eliminate bleed-through and scattering from wire bonds (modified from [28]).

To minimize power consumption, the pixel array is turned off outside the *Imaging* state as described previously. At the beginning of the *Imaging* state, in addition to turning on the pixel array, it is necessary to reset the pixel and drain the previously stored charge on C_int_ with the replica circuit as discussed in [36]. Therefore, with the start of the *Imaging* state, both the pixel array power gating control and the pixel reset control are turned on by the FSM. In cases where the pixel array is not fully switched on during parts of the pixel reset phase, the pixel array settling time affects sensitivity in detecting weak signals from biological samples. In future work, this can be addressed by an earlier start of the pixel array to allow sufficient time for settling before the *Imaging* state. In this work, since the reset timing control is embedded in the FSM, this was only possible by overriding the power gating switch assisted with a wired mode setup while still retaining the power management interface. In future designs, the control over power-gating the imager array will be independently adjustable via the wireless link, ensuring the pixel array is on and in a settled state prior to *Imaging*. For each frame, the serialized pixel data from the on-chip ADC is streamed out to reconstruct the image.

### B. Measurement Results and Analysis

The exposure time of the fluorescence scanner is 2 s and the chip images are taken with T_int_=64 ms. The fluorescence images of the slide samples from untreated and treated mice taken with the scanning microscope and our imager are presented in Fig. 15. CD8 T-cells (shown in pink) are overlaid with the cell nuclei of the entire sample (DAPI, shown in blue). Compared to untreated control samples, the treated mice at later time points are expected to show a significant increase in the population of CD8 T-cells indicating successful immune system activation. The images from our proposed system are consistent with the results of the high-resolution microscope. To quantify the therapeutic response, the T-cell density in the sample is computed by taking the average CD8 T-cell intensity divided by the total nodal area imaged. The comparison between CD8 T-cell density in untreated vs. treated mice is shown in Fig. 16.

**Fig. 15.**
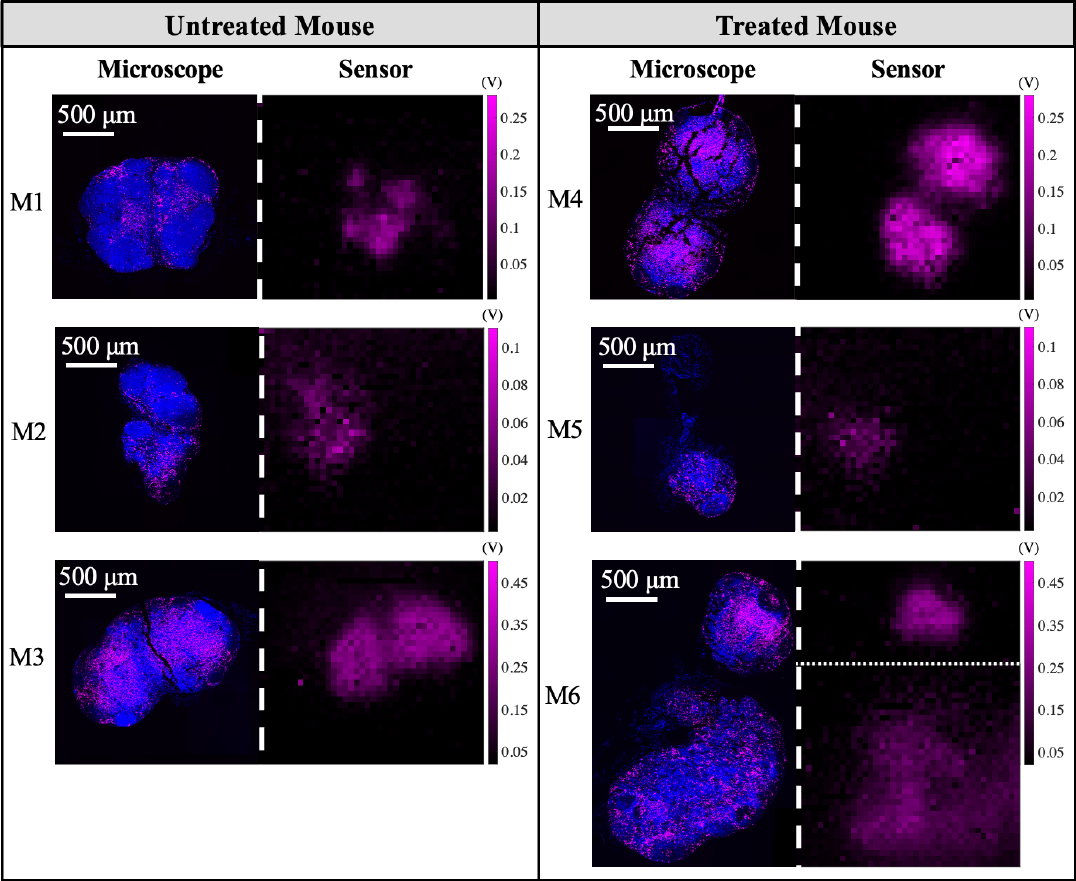
*Ex vivo* images from Zeiss fluorescence slide scanner and our proposed system (T_int_=64 ms). The images are from CD8 T-cells in lymph nodes of mice in an ICI immunotherapy experiment. Images from the 3 untreated mice (M1-M3) are shown in separate rows on the left panel. The images from the mice treated with immunotherapy (M4-M6) are shown on the right. The scale bar is the same for all images. All units for the sensor images are in Volts. The lymph node sample in M6 spans beyond the field of view of the sensor thereby the full image is a composite of 2 overlapping images taken with the sensor).

**Fig. 16.**
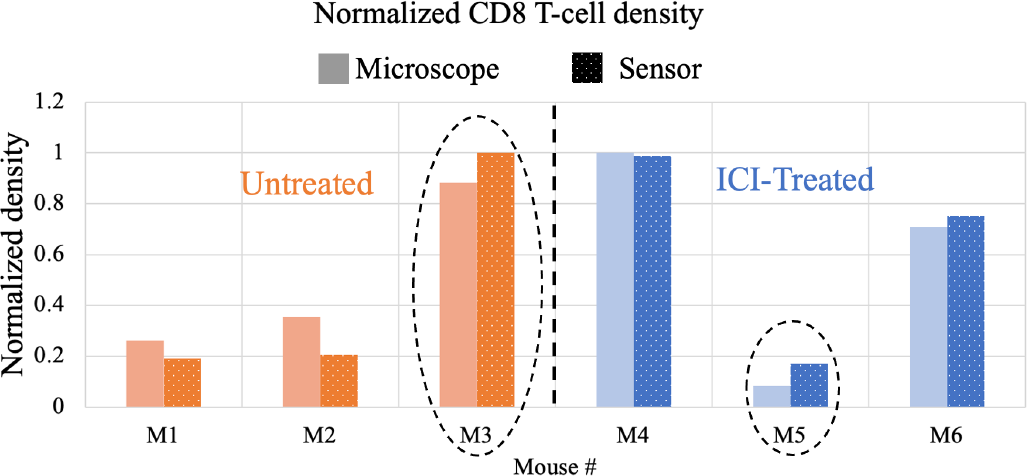
Normalized CD8 T-cell density computed for the sensor and fluorescence microscope images for untreated and treated mice after immunotherapy. The outlier mice samples (M3 and M5) are circled.

On average, CD8 T-cell density increases by 9.8% in the microscope images and 17.2% for the sensor images after immunotherapy, however, owing to mouse-to-mouse heterogeneity, there is wide variability between the baseline of the immune response in different samples resulting in outliers such as M3 and M5. Thus tracking the dynamic response and the change in cell populations is more informative than capturing a single time point image without information about the pre-therapy baseline. This inherent heterogeneity motivates future work for *in vivo* experiments where dynamic changes in response can be observed through implantation of the device without the need for sacrificing the mice and losing the continuous-time data.

## VI. Wireless Mode Measurement Results

After verifying the performance of the individual blocks including the imager, the overall operation of the system is tested by imaging a fluorescent dye (Cyanine5.5-NHS), distributed underneath a standard resolution test target (USAF, Thorlabs). In this measurement, the acoustic interrogation from the transducer powers up the device to capture an image, and then the backscattered pixel data is transmitted back to the transducer to be processed for image reconstruction.

### A. Experimental Setup

Fig. 17 represents the measurement setup. The piezoceramic is placed at a depth of 2 cm away from the US transducer in oil. An acoustic absorber (Aptflex F28P, Precision Acoustics) is used to minimize the reflection from the bottom and sides of the tank. The transducer is controlled by a high-voltage pulser board (Max14808, Maxim Integrated) which is digitally controlled by an FPGA (Opal Kelly XEM6010) to apply the desired interrogation sequence to the chip as previously presented in Fig. 10 (V_piezo+_). Externally, the piezoceramic is connected to the chip in the optical setup where it is fabricated with the optical filter (ET FITC-Cy5, Chroma). The USAF resolution target is covered with Cyanine5.5-NHS (Excitation: 683 nm, Emission: 703 nm) underneath and is positioned on top of the imager array to evaluate the image resolution.

**Fig. 17.**
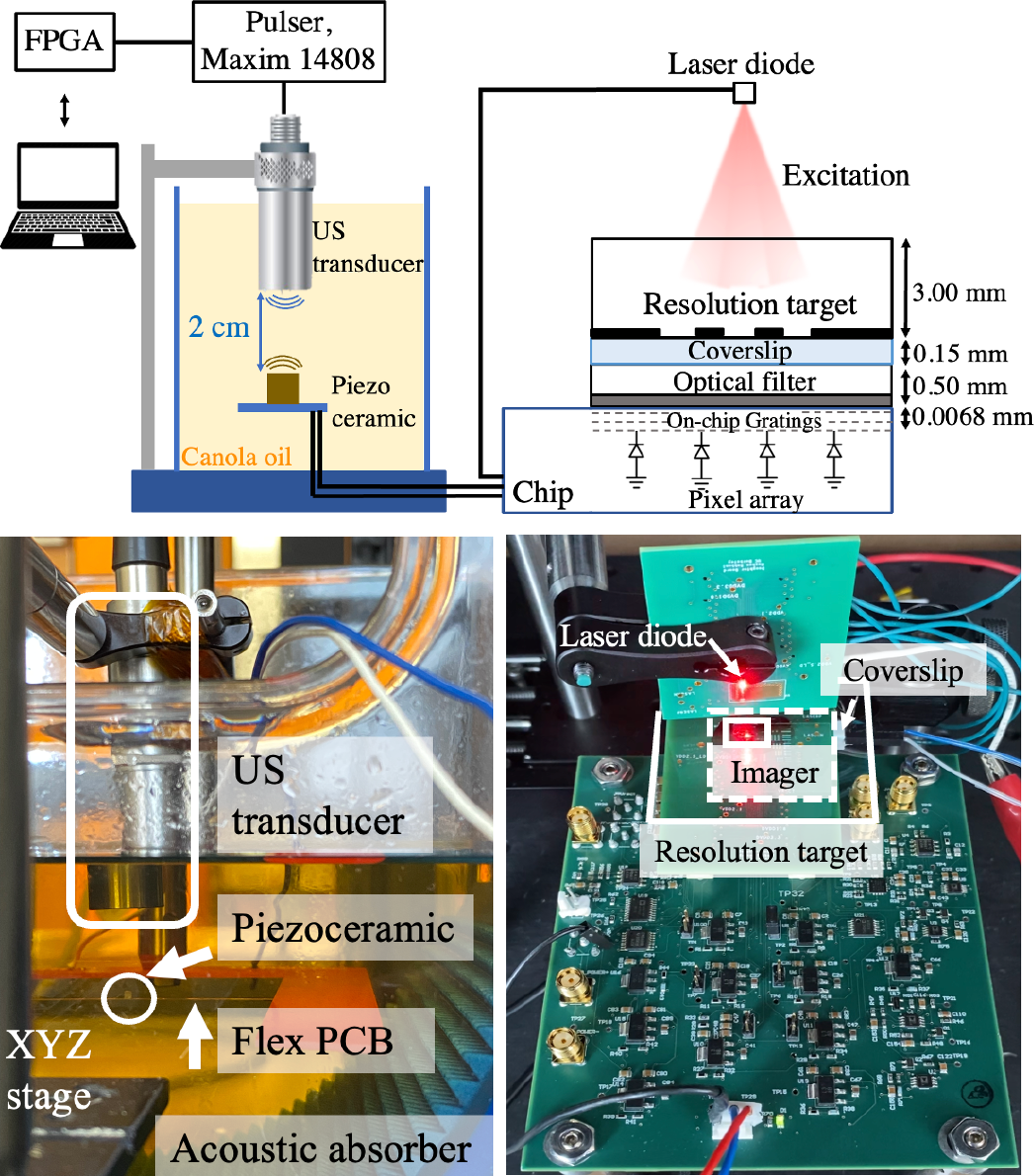
Measurement setup for imaging patterns on a USAF resolution target covered with a coverslip containing Cy5.5 fluorescent dye. On the left, the acoustic setup is shown inside a tank of canola oil. The image on the right is a snapshot of the instant the laser diode is turned on. For visibility purposes, a positive USAF resolution target is chosen over a negative pattern to show the components underneath the target.

### B. Backscattered Images

The backscattered images taken from the highlighted regions on the resolution target are shown in Fig. 18. The images are taken after a 150 s *Charge-Up*. Instead of a continuous waveform, the ultrasound interrogation is 40% duty-cycled to ensure the safe operation of the transducer without being overheated. The images are taken after a T_int_=32 ms and the backscattered data after a total readout time of 389 ms is captured by the transducer. The frame time is sufficient to capture the movements of cells inside the body [47], [48]. Our platform is successful in distinguishing metallic patterns, and features as small as 140 μm with a contrast higher than 87%, making it a viable solution for the detection of clusters of a few hundred cells in immunotherapy. Contrast is calculated from (*V*_*max*_ − *V*_*min*_)/ (*V*_*max*_ + *V*_*min*_ − 2*V*_*bk*_), where *V*_*max*_ and *V*_*min*_ are the values of the bright and dark pixels in a row scan inside the region of interest and *V*_*bk*_ is the background signal. The outlier pixels in the dark region correspond to the BER while backscattering with the lower V_rect_ values. Taking multiple images and averaging can further improve image quality. The details of the post-processing algorithm to remove the outliers are included in Supplementary Fig. 1.

**Fig. 18.**
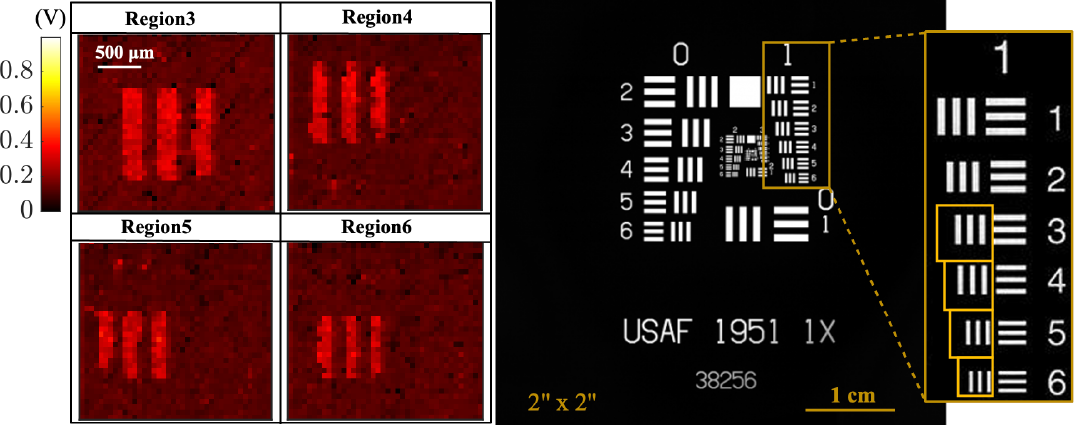
Backscattered images from the highlighted regions on the USAF resolution target with T_int_ = 32ms. The scale bar is in Volts.

## VII. Conclusion

A comparison with recently published implantable imagers is shown in Table I. The wired implantable imagers [22], [24], [25] perform high-resolution fluorescence microscopy using CMOS photodiodes or single photon avalanche diodes (SPADs). Their utility for continuous monitoring is restricted by the risk of infection due to external wiring. While the thermoacoustic sensor [26] provides a wireless interface for untethered implantation, multicellular-level detection for biological applications is not feasible without enhancing the resolution.

**TABLE I.**
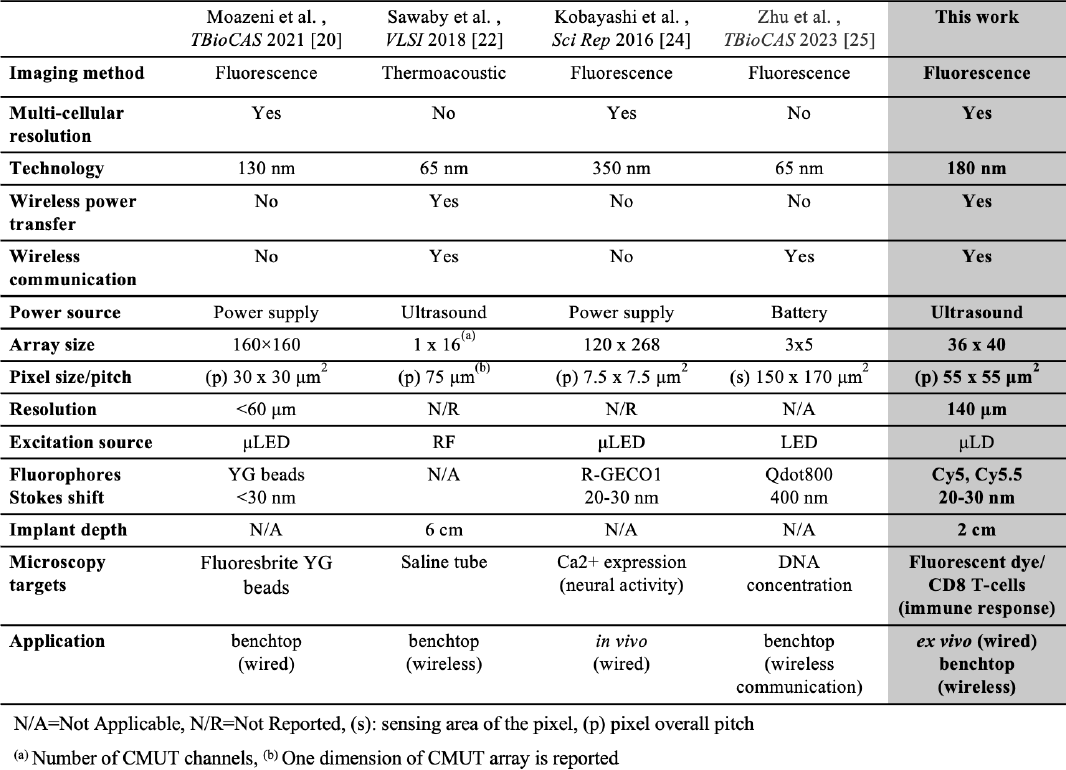
Comparison Table of Implantable Image Sensor

We demonstrate, to the best of our knowledge, the first platform for wireless untethered chip-scale fluorescence microscopy with the systematic device-level integration of the optical source eliminating bulky optics such as focusing lenses and fibers and electrical components such as batteries or any external wiring. We propose a proof-of-concept prototype for real-time microscopy, within the tissue, at early time points of therapy, to address a key challenge in assessing therapeutic response and disease progression. The system utilizes the stored wireless energy from an ultrasound link to provide the high 78 mW instantaneous power required for the optical source in fluorescence imaging during a 32 ms integration time. For each frame, 11.52 kbits of image data are transmitted via ultrasound backscattering using the same piezoceramic transceiver used for power transfer. The imager array can capture high-resolution *ex vivo* images of CD8 T-cell profiles consistent with images taken with a high-resolution fluorescence microscope. The overall system performance is demonstrated by resolving 140 μm features on a USAF resolution test target and transmitting the data wirelessly to the external US transducer.

Future work to further enhance sensitivity, integrate the system in a biocompatible structure and minimize the form factor (summarized in the Supplementary) will enable the utility of the device for real-time fluorescence microscopy to detect microscopic cell foci, increasing visibility into the tumor microenvironment and guiding cancer therapy.

## Acknowledgment

The authors would like to thank sponsors of BSAC (Berkeley Sensors and Actuators Center) and TSMC for chip fabrication, the UCSF Preclinical Therapeutics Core, UCSF Histology and Biomarker Core, and Hui Zhang for mouse experiment preparation, and Mauricio Bustamante for piezoceramic assembly.

## Notes

1 This work was supported by the Office of the Director and the National Institute of Dental & Craniofacial Research of the National Institutes of Health under Award DP2DE030713

### Competing Interest Statement

The authors have declared no competing interest.

