## Supplementary Material for "Towards A Wireless Image Sensor for Real-Time Fluorescence Microscopy in Cancer Therapy"

Rozhan Rabbani, *Member, IEEE,* Hossein Najafiaghdam*, Member, IEEE,*
Micah Roschelle*, Member, IEEE,* Efthymios Philip Papageorgiou, *Member, IEEE,*
Biqi Rebekah Zhao*, Member, IEEE,* Mohammad Meraj Ghanbari, *Member, IEEE,*
Rikky Muller, *Senior Member, IEEE,* Vladimir Stojanovic, *Senior Member, IEEE,*
*and* Mekhail Anwar, *Member, IEEE*

#### Comparison of the current illumination scheme with implanted setup

The implanted setup as shown in the conceptual diagram in Fig. 1(a) requires the laser diode to be assembled next to the sensor while illuminating the target via epi-illumination. Compared to trans-illumination in the current setup shown in Fig. 1(b) and Fig. 17, epi-illumination lowers the background signal due to the excitation light being reflected off the surface of the sample and not directly incident on the surface of the imager. This can positively affect the signal-to-background ratio. However, there are additional effects on signal intensity that need to be addressed:

1. **Spacer thickness:** To deliver light via epi-illumination from the edge emitter laser diode to the sample, a glass spacer between the sensor and the target is required. The thickness of the spacer increases the distance between the source and the target lowering the light intensity absorbed by the fluorophores.
   This effect can be studied using the simplified illumination models shown below:

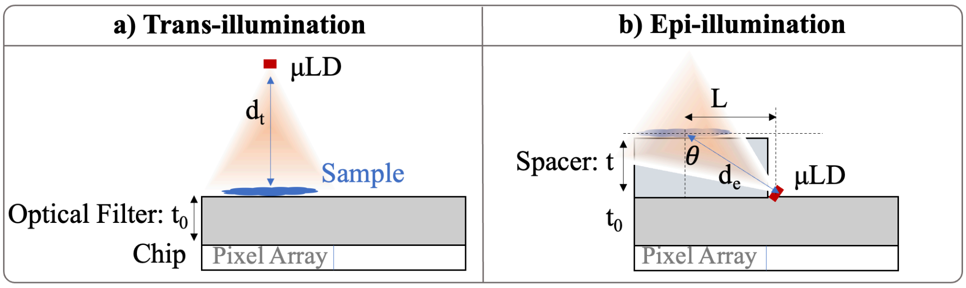


**Supplementary Figure 1.** Trans-illumination and epi-illumination setups with the chip, optical filter, µLD and glass spacer. The pixel array covers about 40% of the entire chip area. The spacer length (L) is chosen to be the same as the pixel array length (2.2mm). The µLD is placed as close as possible to the spacer.

The intensity of the light received incident on the same surface area of the sample in both cases can be calculated as shown below:

$$E = \frac{I}{d^2}$$

E is the irradiance at distance d from a point source of light with an overall intensity of I

$$P_{sample,trans} = E_{trans} A_{Sample}$$

Where $P_{sample,trans}$ is the light intensity received by a surface area of $A_{Sample}$ being trans-illuminated from a distance of $d_t$

$$P_{sample,epi} = E_{epi} \cos\theta \, A_{Sample}$$

Where $P_{sample,epi}$ is the light intensity received by a surface area of $A_{Sample}$ from epi-illumination at a distance of $d_e$ and incident angle of $\theta$.

The irradiance at the sensor surface from the fluorescent sample is proportional to:

$$E_{sensor,trans} \propto P_{sample,trans} \frac{1}{(t_0)^2}$$
$$E_{sensor,epi} \propto P_{sample,epi} \frac{1}{(t + t_0)^2}$$
Where $t$ is the thickness of the glass spacer and $t_0$ is the thickness of the optical filter.

2. **Reflection due to oblique incidence:** Compared to normal incidence, oblique illumination increases reflection at the interface between air and the spacer reducing the transmitted light to the target.

The reflection of light at the intersection of air and a second medium with a refractive index of n can be calculated from:
$$R_p = \left(\frac{\cos\phi - n\cos\,\alpha}{\cos\phi + n\cos\,\alpha}\right)^2, R_s = \left(\frac{\cos\alpha - n\cos\,\phi}{\cos\alpha + n\cos\,\phi}\right)^2$$
Where $R_p$ and $R_s$ refer to reflections of TM and TE waves, respectively [1]. $\alpha$ is the angle of the incoming beam in air and $\phi$ is the angle of the transmitted rays in the second medium. For normal incidence reflection can be simplified to
$$R = \left(\frac{1-n}{1+n}\right)^2$$
The ratio of light transmitted at the intersection of air and the medium n can be calculated from:
$$T = 1 - R$$
Assuming n=1.45 for tissue, 96.6% of the light will reach the sample in Suppl. Fig.1 (a). For oblique incidence, the transmitted power from air to glass (n=1.5) for a range of incoming angles is shown in Suppl. Fig. 2(a). The reflection coefficient is not calculated for the interface of glass-tissue because of their similar refractive indices.
Combing the effect of distance (part 1) and reflection (part 2) the relative irradiance of the emitted light for the same laser power is plotted as a function of the thickness of the spacer in Suppl. Fig. 2(b). The process is repeated for a range of µLD-sample distances in trans-illumination. The plots are generated considering a 500µm thick optical filter, s-polarized light with lower transmission for worse case, and L=2.2mm in Suppl. Fig. 1 (b). The dashed lines are generated considering the effect of reflection for both trans-illumination and epi-illumination.

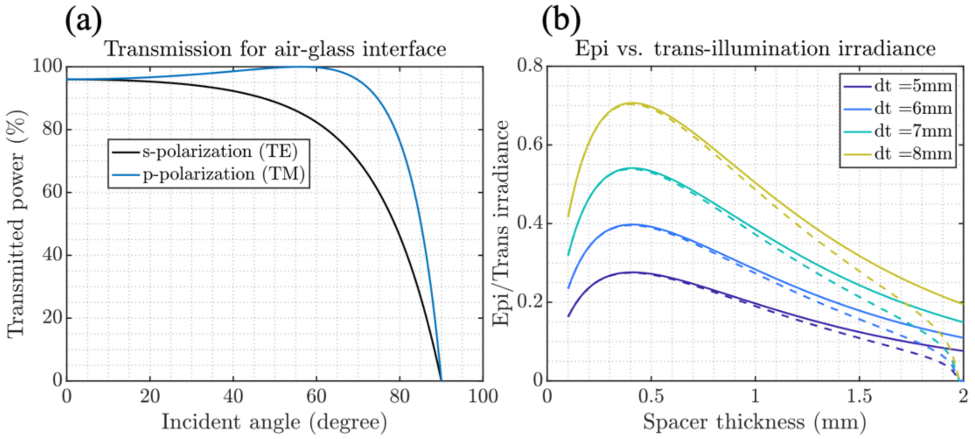


**Supplementary Figure 2. (a)** Transmission of light from air to glass for different incoming angles of incident. (b) comparison of the irradiance of

As shown in Suppl. Fig. 2(b), compared to trans-illumination at distance of $d_t$=7mm (similar to the experimental setup), with an optimized spacer width of 400µm, the intensity is reduced by 45% for epi-illumination. The loss in signal intensity can be improved by increasing the integration time for each frame. Another effect is the lower resolution due to the larger distance of the sample from the sensor caused by the spacer which linearly diminishes resolution [2].

3. **Nonuniform illumination:** One solution to improve uniformity is using light-guide plates (LGP) [3], [4] to deliver light from the laser to the tissue and improve uniformity of the light profile within the sensor field of view. Without LGPs, nonuniform illumination can be characterized and corrected computationally [5]. In [6], the authors use convolutional neural networks to generate a uniform image from the images taken under nonuniform illumination conditions. The work in [7] summarizes mathematical approaches that are used to correct for uneven illumination in digital images.
   Given these methods the following steps are needed to correct images based on the nonuniform illumination profile by capturing an illumination map:
   a. The variability of the pixel responsivity can be captured by taking an image of a fluorescent dye, spread evenly on a glass slide covering the imager array, illuminated by a wide collimated laser beam from the top.
   b. Once the pixel-to-pixel variation map is determined, the laser diode in the implanted setup can turn on to illuminate the uniformly distributed dye on the sensor. Using (a) and (b) an image of the illumination profile can be obtained.
   c. This nonuniform illumination map can be used with machine learning-based methods or computational algorithms to correct future captured images for the effects of non-uniform illumination.

#### Illumination and optical power safety requirements

The laser diode used in this work is a class III medical laser ($P_{out}$< 5 mW). According to the American National Standard for Safe Use of Lasers (ANSI Z136.1-2014) the maximum exposure equal to $1.1t^{0.25}$ J/cm$^2$. Where t refers to the total exposure time of the laser. For an integration time of $T_{int}$=64 ms, the maximum radiant exposure allowed is 0.55 J/cm$^2$. With the current optical output power, the radiant exposure $H_{e,}$ is 50 mW/cm$^2$ * 64 ms= 0.0032 J/cm$^2$ which is more than 170x lower than the ANSI limit.

#### Image outlier detection and correction

In wireless measurements, the bit error rate from backscattering can lead to outlier pixels in the reconstructed images. The outliers can be detected with an algorithm that compares the value of each pixel with the surrounding pixels. For each pixel, the mean and standard deviation of the 8 neighboring pixels is computed (except for the edge or corner pixels with 5 and 3 neighboring pixels respectively). Once the pixel value falls outside a certain range according to the statistics of the surrounding samples ($\mu\pm2\sigma$ in this case, where $\mu$ is the mean and $\sigma$ is the standard deviation of the neighboring pixels excluding the pixel of interest), its value is replaced by the average of the surrounding pixels. The algorithm is built on the function proposed here. The images before and after applying the outlier detection are shown below:

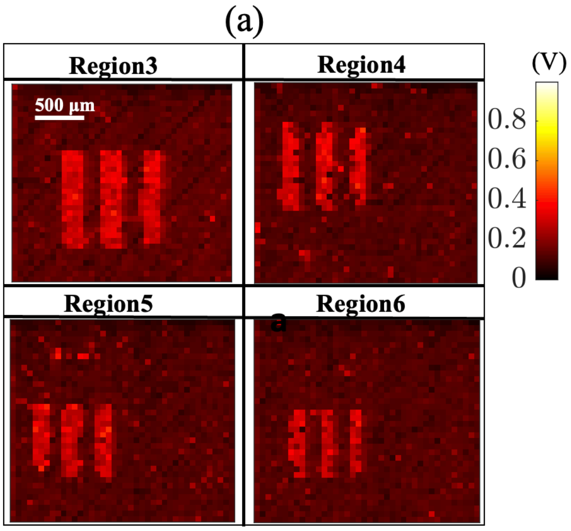


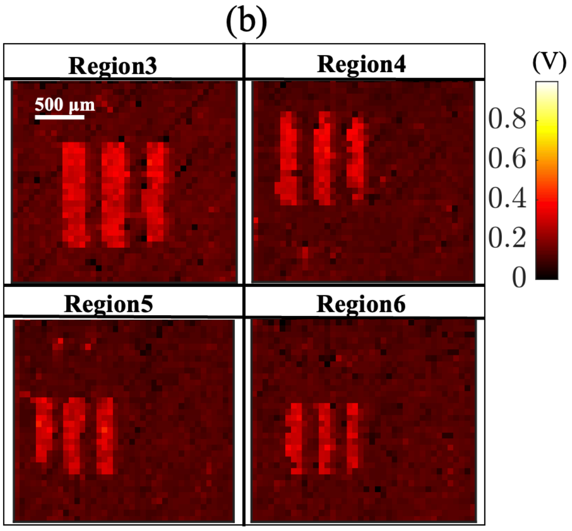

**Supplementary Figure 3.** Outlier detection and correction for backscattered images. (a) The initial backscattered images of the highlighted regions in the USAF resolution target in Fig. 19. **(b)** Same image using the outlier removal algorithm based on the mean and standard deviation of the neighboring pixels. The units of pixels are in Volts and the scale bar is the same for all images.

### Future directions for *in vivo* imaging

The proposed device is a proof-of-concept wireless fluorescence image sensor that exhibits fluorescence microscopy with sufficient resolution unlocking the potential for untethered imaging inside the tissue. Here is a summary of the key steps to facilitate the device's utility for *in vivo* applications:

- Laser diode

  Use of a laser diode with higher electrical to optical efficiency (such as CHIP-650-P5, Roithner Laser Technik with a 12.5% efficiency compared to the current diode with 4.4% efficiency) lowers the required stored charge in $C_{store}$. This leads to a more than 64% reduction in the size of $C_{store}$ while maintaining the same optical power.
- $T_{int}$ reduction and averaging

  The size of the storage capacitor is chosen to be 0.8 mF for this design. For the shot noise limited region of operation in Fig. 5, averaging two consecutive images each taken with half of the standard $T_{in}$ maintains the SNR within 1dB of the original value. This can lower the size of $C_{store}$ by 50%.
- Imager array

  The scalable design of the CMOS imager sensor allows various array sizes tailored to the physical requirements of the *in vivo* application.
- Sensitivity enhancement

  The imager array's power-gating control will be adjustable independently in the future, which would guarantee that the imager is turned on and settled before starting the *Imaging* state. This will improve the sensitivity of the imager to detect weak biological signals on the order of the photodiode dark current (~14 aA/μm$^2$).
